# RORA regulates neutrophil migration and activation in zebrafish

**DOI:** 10.1101/2021.12.03.470833

**Authors:** Alan Y. Hsu, Tianqi Wang, Ramizah Syahirah, Sheng Liu, Kailing Li, Weiwei Zhang, Jiao Wang, Ziming Cao, Simon Tian, Sandro Matosevic, Chris Staiger, Jun Wan, Qing Deng

## Abstract

Neutrophil migration and activation are essential for defense against pathogens. However, this process may also lead to collateral tissue injury. We used microRNA overexpression as a platform and discovered protein-coding genes that regulate neutrophil migration. Here we show that miR-99 decreased the chemotaxis of zebrafish neutrophils and human neutrophil-like cells. In zebrafish neutrophils, miR-99 directly targets the transcriptional factor *RAR-related orphan receptor alpha (roraa)*. Inhibiting RORα, but not the closely related RORγ, reduced chemotaxis of zebrafish and primary human neutrophils without causing cell death, and increased susceptibility of zebrafish to bacterial infection. Expressing a dominant-negative form of Rorα or disrupting the *roraa* locus specifically in zebrafish neutrophils reduced cell migration. At the transcriptional level, RORα regulates transmembrane signaling receptor activity and protein phosphorylation pathways. Our results, therefore, reveal previously unknown functions of miR- 99 and RORα in regulating neutrophil migration and anti-microbial defense.

## Introduction

Inflammation is essential for the restoration of tissue homeostasis after injury and infection. Neutrophils, the primary effector cells of acute inflammation, are the first to infiltrate inflammation sites. They combat disease by phagocytosing foreign particles and secreting cytokines, anti-microbial molecules, and chromatin while contributing to collateral tissue injury [1]. Under inflammatory conditions, neutrophils are a heterogeneous population with pro-and anti-inflammatory phenotypes that cooperate to resolve inflammation [2; 3]. In line with their multifaceted roles, neutrophils contribute to dangerous, chronic inflammatory conditions in humans, including those associated with tumors, chronic obstructive pulmonary disease, arthritis, and adipose inflammation. On the other hand, neutrophil activity is beneficial to wound healing, infarction repair, and inflammatory bowel diseases [4]. Enhancing, inhibiting, or restoring neutrophil function are attractive therapeutic strategies [5]. In clinical trials, there are neutrophil-focused drug candidates, including CXCR2 blockades, neutrophil elastase inhibitors, and C5a receptor antagonists. Nanotechnology-based drug delivery [6] has shown promise in blocking the airway and systemic inflammation [7]. As vehicles that efficiently cross the blood vessel barrier, neutrophils also provide a novel way to deliver therapeutic cargos to inflammation sites [8]. A better understanding of the neutrophil-intrinsic mechanisms that regulate neutrophil migration will have broad translational importance in preventing and treating a wide range of inflammation-related diseases.

MicroRNAs are small RNA molecules of 22-24 nt that regulate homeostasis in health and disease [9]. MicroRNAs primarily suppress target gene expression by binding microRNA seed sequences (nucleotides 2-8) to partially complementary sequences in their transcripts’ 3’ untranslated regions. MicroRNA “mimics” and “inhibitors” are emerging as next-generation therapeutics because of their ability to modulate a network of genes [10; 11]. The first small RNA drug, ONPATTRO, a small-interfering RNA, is approved for treating a rare polyneuropathy [12]. Candidate microRNA therapeutics are in clinical development for other human conditions such as HCV infection [13], wound healing defects and myocardial infarction [14], and cancer [15]. In addition, microRNAs are being used as tools to discover novel regulators of biological processes.

There are challenges associated with studying terminally differentiated neutrophils; namely, only a limited set of genetic tools are available to examine the plasticity of the cells *in vivo*. To address this challenge, we are using zebrafish, a genetically tractable vertebrate model with a well-conserved innate immune system [16]. Adaptive immunity is morphologically and functionally mature only after 4-6 weeks postfertilization. Therefore zebrafish embryonic provides a unique tool to investigate innate immunity without the impact from the adaptive immune system [17].

Through a phenotypic screen in zebrafish, we identified nine microRNAs that, when overexpressed, suppressed neutrophil migration [18]. Among the positive hits, dre-miR-99-1 significantly reduced neutrophil chemotaxis. The current study characterized miR-99 substrates and discovered an unexpected role of the RAR-related orphan receptor (RORα) in regulating neutrophil migration.

## Materials and Methods

### Generation of transgenic zebrafish lines

The zebrafish experiment was conducted in accordance with internationally accepted standards. The Animal Care and Use Protocol was approved by The Purdue Animal Care and Use Committee (PACUC), adhering to the Guidelines for the use of Zebrafish in the NIH Intramural Research Program (protocol number: 1401001018). Zebrafish *roraa* gene was cloned from zebrafish mRNA using SuperScript III RT-kit (Invitrogen #18080044) and amplified with the following primers. zroraa wt+: 5′- aaaaccccggtcctatgcatatgatgtattttgtgatttcagctatgaaagct-3′; zroraa wt-: 5′-ctgattatgatctagactacccgtcaacgggcat-3; zroraa LBD deletion DN+: 5′- aaaaccccggtcctatgcatatgatgtattttgtgatttcagctatgaaagctcaaatcg-3′; zroraa LBD deletion DN -: 5′- ctgattatgatctagagtccaggccggattgatcagg-3 and inserted into a Tol2-lyzC-mcherry-2A backbone by using infusion cloning kit (Takara #638920). The construction method for neutrophil-specific Cas9 expression and the guide RNA expression fish lines has been described in our previous study [40]. The protospacer sequences for targeting *dre*-*roraa* in this research are 5′- taattccctgcaagatctg-3′; 5′-gcacgttatcacgccataa-3′, both guide RNAs target on *dre roraa* exon2. Microinjections of fish embryos were performed by injecting 1nl of a mixture containing 25 ng/µl plasmids and 35 ng/µl tol2 transposase mRNA in an isotonic solution into the cytoplasm of embryos at the one-cell stage. The stable lines were generated as previously described [19]. At least two founders (F0) for each *line* were obtained. Experiments were performed using F2 larvae produced by F1 fish derived from multiple founders to minimize the artifacts associated with random insertion sites.

### RNA seq

Kidney marrow was dissected from 3 adults from *Tg(lyzC: miR-99-1-dendra2)^pu26^*, *Tg(lyzC: vector-dendra2)^pu7^*, *Tg(lyzC: mcherry-2a) ^pu29^* and *Tg(mcherry-2a-*RORα DN*) ^pu30^* and neutrophils were sorted using FACS. Total RNA was extracted using RNeasy Plus Mini Kit (Qiagen #74104). RNAseq was performed at The Center for Medical Genomics at Indiana University School of Medicine. Samples were polyA enriched and sequenced with Illumina HiSeq 4000 ultra-low with reads range from 37M to 44M. The RNA-seq aligner from the STAR (v2.5) [20] was employed to map RNA-seq reads to the reference genome, zebrafish (GRCz11), with previously the following parameter: “--outSAMmapqUnique 60”. Uniquely mapped sequencing reads were assigned to genes using featureCounts (from subread v1.5.1) [21] with the following parameters: “-p -Q 10”. The genes were filtered for further analysis if their count per million (CPM) of reads was less than 0.5 in more than three samples. The method of a trimmed mean of M values (TMM) was adopted for gene expression normalization across all samples, followed by differential expression analysis between different conditions using edgeR (v3.20.8). Differentially expressed gene was determined for the comparison if its false discovery rate

(FDR) adjusted p-value was less than 0.05 and the amplitude of fold change (FC) was larger than linear 2-fold. The functional analysis was performed on DEGs of our interest with a cutoff of FDR < 0.05 to identify significantly over-represented Gene Ontology (GO) and/or KEGG pathways by using the DAVID [22]. To compare the down-regulated DEG set in rorα DN neutrophil to the published THP-1/HUVEC RORα CHIP-seq data, the genes from CHIP-seq result were converted to their orthologues in zebrafish. The gene duplicates after the conversion from human cells were removed. The zebrafish genes not having the orthologues to human were also removed before comparison.

### Quantitative RT-PCR

Total RNA was purified using MiRVANA miRNA purification kit (ThermoFisher) for miRNA or RNeasy Plus Mini Kit (Qiagen #74104) for mRNA. MicroRNAs were reverse transcribed with Universal cDNA Synthesis Kit II (Exiqon). MicroRNA RT-qPCR was performed with ExiLENT SYBR® Green master mix (Exiqon), and mRNA with one-step RT-qPCR was performed with SuperScript® III Platinum® SYBR® Green kit (Invitrogen) using LightCycler ® 96 Real-Time PCR System (Roche Life Science). The specificity of the primers was verified with a single peak in the melt curve. The relative fold change with correction of the primer efficiencies was calculated following instructions provided by Real-time PCR Miner (http://ewindup.info/miner/data_submit.htm) and normalized to *U6*. miRNA Primers used in this study were purchased from Exiqon. The relative fold change with correction of the primer efficiencies was calculated following instructions provided by Real-time PCR Miner and normalized to *rpl32*. Primers: dre-grn1+: 5′-aacccagccagcaagatg-3′, dre-grn1−: 5′- ccaccgggatagacagatca-3′, dre-fabp7a+: 5′-cagcagacgatagacatgtgaagt-3′, dre-fabp7a−: 5′- tcccacctctgaacttggac-3′, dre-il1b+: 5′-atgctcatggcgaacgtc-3′, dre-il1b−: 5′-tggttttattgtaagacggcact-3′, dre-tgfbr1+: 5′-tgcaacaagaacccaaaagtt-3′, dre-tgfbr1−: 5′-gggactcatagactggggttc-3′, dre-roraa+: 5′-gaatgatcagatagtgcttctcaaag-3′, dre-roraa−: 5′-cacggcacattctgacga-3′, dre-rorc+: 5′- tcttttcctatccaacctctctaca-3′, dre-rorc−: 5′-gagtggtctctttatgtgagcgta-3′, dre-rpl32+: 5’- tcagtctgaccgctatgtcaa-3’; dre-rpl32-: 5’- tgcgcactctgttgtcaatac-3’; dre-ef1a+:5’- tgccttcgtcccaatttcag-3’; dre-ef1a-:5’- taccctccttgcgctcaatc-3’; dre-roraa-overexpression+:5’- tcaggcatccattatggcg-3’; dre-roraa-overexpression-:5’- cggtcaatcaggcagttctt-3’; dre-arhgap17b+:5’- ttggcggatgaagaggattc-3’; dre-arhgap17b-:5’- ccaatcgcacattttccagg-3’; dre- ptk2ba+:5’- attacatgcagcacaacgcc-3’; dre-ptk2ba-:5’-ccagacccacatctttctcca-3’; dre-mhc1zba+:5’- gaagattcccaaacagcactgg-3’; dre-mhc1zba-:5’-cgttccatcagaatgttgacgttc-3’; dre-tcerg1l+:5’- aagcagagaacagcggataag-3’; dre-tcerg1l-:5’-ttagcctctgggtttggattg-3’.

### Dual-luciferase reporter assay

zebrafish *roraa* 3′UTR was amplified from zebrafish genomic DNA with SuperScript III RT-kit (Invitrogen #18080044) and cloned into psiCHECK2 (Promega) at *Xho*I and *Not*I cloning sites with the following primers: zroraa+:5’- taggcgatcgctcgacaaggcagctcactaggaacagaactg-3’; zroraa-: 5’- ttgcggccagcggccggaggacagggaaacgtagctgtac-3’. Mutated 3’UTR constructs were generated using Infusion HD cloning kit (Clontech) with the following primers: zroraa mut+: 5’- acaaaatgcccattctacagggtaaccactgctgtgg-3’; zroraa mut-: 5’- gaatgggcattttgtcgagcgacggacgtc-3’.

DNA encoding *miR-99* or vector was amplified from the construct used for expression in zebrafish and inserted into pcDNA3.1 at the *Hin*dIII/*Xba*I cloning sites using the following primers: pcDNA+: 5;- gtttaaacttaagcttgccaccatggatgaggaaatcgc-3’; pcDNA-: 5’- aaacgggccctctagagaccggtacccccgggctgc-3’.

### Live imaging

Larvae at three dpf were settled on a glass-bottom dish, and imaging was performed at 28 °C. Time-lapse spinning disk confocal microscopy (SDCM) was performed with a Yokogawa scanning unit (CSU-X1-A1) mounted on an Olympus IX-83 microscope, equipped with a 20X 0.5–numerical aperture (NA) UPlanSApo oil objective (Olympus) and an Andor iXon Ultra 897BV EMCCD camera (Andor Technology). GFP and mCherry were excited with 488 nm and 561 nm and fluorescence emission collected through 525/30-nm and 607/36-nm filters, respectively, to determine the localization of stable actin in migrating neutrophils in vivo. Images were captured using MetaMorph version 7.8.8.0 software at 10 s intervals of a total of 5 min.

Time-lapse fluorescence images in the head mesenchyme were acquired with a laser-scanning confocal microscope (LSM710, Zeiss) with a Plan-Apochromat 20×/0.8 M27 objective. The fluorescent stacks were flattened using the maximum intensity projection and overlaid with a single slice of the bright-field image. The velocity of neutrophils was quantified using ImageJ with the MTrackJ plugin.

### Inflammation assays in zebrafish

Zebrafish wounding and infection was performed as described [23]. Briefly, three dpf larvae were amputated posterior to the notochord or inoculated with *P. aeruginosa* (PAK) into the left otic vesicle or the vasculature at 1000 CFU/embryo. The larvae were fixed in 4% paraformaldehyde at one h post-wounding or -infection. Neutrophils were stained with Sudan Black, and the number at the indicated regions was quantified.

### Generation of stable HL-60 cell lines

HL-60 cells were obtained from ATCC (CCL-240) and cultured using RPMI-1640 with HEPES supplemented with 10% FBS and sodium bicarbonate. The lentiviral backbone pLIX_403 was a gift from David Root (Addgene plasmid # 41395). The DNA sequence flanking *mir-99* was cloned from HL-60 cell genomic DNA and cloned into a backbone containing *dendra2*. The miRNA and *dendra2* reporter were then cloned into pLIX_403 vector using the NheI/AgeI sites with primer set: pLIX-mir+: 5’ tggagaattggctagcgccaccatggatgaggaaatcgc-3’, pLIX-mir-: 5’- catacggataaccggttaccacacctggctgggc-3’. Stable HL-60 cell lines were generated as described [24; 25]. Briefly, HEK-293 cells were transfected with pLIX_403 or pLKO.1, together with VSV-G and CMV8.2 using lipofectamine 3000 (ThermoFisher). The viral supernatant was harvested on day three and concentrated with Lenti-X concentrator (Clontech 631232). HL-60 cells were infected with concentrated lentivirus in complete RPMI medium supplemented with four µg/ml polybrene (Sigma TR-1003-G) at 2500g for 1.5h at 32 °C and then selected with one µg/ml puromycin (Gibco A1113803) to generate stable lines.

### Flow cytometry

Cells were stained with Alexa Fluor 647 conjugated CD11b (neutrophil differentiation marker, Biolegend, 301319, clone IV-M047, 2ul to 1 x 10^6^ cells) or the isotype control antibody (Biolegend, 400130, clone MOPC-21, 2ul to 1 x 10^6^ cells), RUO AnnexinV (apoptosis marker, BD, 563973, 5ul to 1 x 10^6^ cells) and RUO propidium iodide (dead cell marker, BD, 556463, 5ul to 1 x 10^6^ cells). Cells were stained in staining buffer (1% BSA, 0.1% NaN3) and incubated on ice for 1 hour, washed three times with staining buffer, and resuspended in suitable volumes.

Following the manufacturer’s protocol, cell cycle profiling was obtained using Vybrant™ DyeCycle™ Ruby Stain (Thermo #V10309). Fluorescence intensity was collected using BD LSR Fortessa. Results were analyzed with Beckman Kaluza 2.1 software.

### Transwell migration assay

Transwell assays were performed as described [24]. Briefly, 2x10^6^ cells/ml differentiated HL-60 cells were resuspended in HBSS with 0.5%FBS and 20 mM HEPES. 100 µl were placed into a 6.5 mm diameter 3 µm pore size transwell insert (Corning#3415) and allowed to migrate for two h at 37°C towards 100 nM fMLP in 500 µl of HBSS in a 24-well plate. Loading controls were done by directly adding 100 µl of cells to 400 µl of HBSS. Cells that migrated to the lower chamber were released with 0.5 M EDTA and counted using a BD LSRFortessa flow cytometer with an acquisition time of 30 s. The counts were normalized with the total numbers of cells added to each well the data was then gated for live cells and analyzed with Beckman Kaluza 2.1 software.

### NETosis assay

Differentiated HL-60 cells (dHL60) were resuspended in HBSS in 20 mM HEPES with 0.5% FBS (mHBSS) and allowed to attach to fibrinogen-coated slides for 30 min at 37°C. Neutrophil extracellular traps (NETs) were induced with 50 nM PMA (sigma #P1585) in HBSS for 4 h at 37°C. NETs were enumerated using cell-permeable Hoechst 33258 at 1 µg/ml (Thermo #62249) and cell impermeable Sytox Green at 1 µM (Thermo #S7020). Images were acquired using Lionheart FX Automated Microscope (Biotek) 10x phase objective, Plan Fluorite WD 10 NA 0.3 (1320516). The percentage of cells forming NETs was calculated by dividing the number of Sytox Green-positive cells by that of the Hoechst-positive cells.

### ROS assay

Amplex™ Red Hydrogen Peroxide/Peroxidase Assay Kit (Thermo #A22188) was used to detect extracellular ROS. dHL60 cells (20,000 cells per well) were incubated with 0.1µg/mL of PMA (sigma #P1585) in 96 well black/clear bottom plate (Thermo #165305). ROS reading was taken 30 - 45 minutes after stimulation using Synergy Neo2 (BioTek) at Ex/Em of 571/585 nm. The amount of hydrogen peroxide release was calculated by comparing to standards prepared.

### Phagocytosis assay

BioParticles from pHrodo™ Green BioParticles® Conjugates (Thermo # P35366) were reconstituted and sonicated for 5 minutes at 10% amplitude with 10 seconds intervals. dHL60 cells (5x10^5^) were resuspended in 100uL of BioParticle suspension and incubated at 37°C for 1 hour. The reaction is stopped by placing on ice before running flow cytometry to quantify the mean FITC intensity in the cell population. Cells with beads placed on ice for 1 hour were used as the negative control.

### Inhibitor treatment for zebrafish larvae

ROR inhibitors SR3335 (Cayman # 12072), SR2211 (Cayman # 11972), SR1001 (Cayman # 10922), and VPR-66 (Novus # NBP2-29335) were dissolved in DMSO to make a 100 mM stock, then further diluted in E3 to working concentrations (10-100 µM for recruitment and motility assays, and indicated concentration for survival assays). For neutrophil recruitment and random motility assays, larval were pretreated with the inhibitor for one h before experimental procedures. For survival assays, larval were pretreated and kept in the inhibitors from 1-hour post-infection.

### Primary neutrophil isolation and chemotaxis assay

Primary human neutrophils were obtained from the peripheral blood of healthy adult donors, collected under approval by Purdue University’s Institutional Review Board (IRB). Primary human neutrophils were isolated with Milteny MACSxpress Neutrophil Isolation Kit (Macs # 130-104-434), and RBC lysed for 10 min RT (Thermo # 00-4300-54). 3x10^6^ cells/ml PMNs were incubated in HBSS with 0.1% FBS and 20mM HEPES with or without respected inhibitors for 1 hour. Cells were then mixed 3:1 with matrigel (corning # 356234) and loaded into collagen-coated IBIDI chemotaxis µ-slides (ibidi # 80326) and incubated at 37°C for 20 minutes to allow 3D matrix formation. 15 μl of 1000 nM fMLP was loaded into the right reservoir yielding a final fMLP concentration of 187 nM. Chemotaxis was recorded every 1 min for two h, with a Lionheart FX Automated Microscope (Biotek) 10x phase objective, Plan Fluorite WD 10 NA 0.3 (1320516). Chemotaxis tracking was done with the MTrackJ image J plugin and plotted in Prism 6.0 (GraphPad). Cell viability of primary human neutrophils treated with ROR inhibitor SR3335 (Cayman # 12072) was measured by staining with propidium iodide and analyzed using flow cytometry.

### Mutational efficiency quantification

To determine the mutation efficiency in *Tg(LyzC:Cas9, Cry:GFP, U6a/c: rora guides, LyzC:GFP)*, the *roraa* loci around the sgRNA binding site was amplified by PCR using the following primers: *dre roraa*+, 5′- cgactggtgacagtgaaaaatc-3′; *dre roraa*-, 5′- tttagcttcttaccttgcagcc-3′. PCR products were deep sequenced using WideSeq at the Purdue Genomics Core Facility. Mutation efficiencies were calculated using the CRISPResso2 package (www.github.com/pinellolab/CRISPResso2).

### Survival assay

Larvae at three dpf were injected with one nl of 25 ng/nl LPS or 1000 CFU *P. aeruginosa* (PAK) into the tail vein and incubated individually in 96-well plates. Survival was tracked for four days or when one group reached 100% mortality. Representative results of at least three independent experiments (n≥20 larvae in each experiment) were shown.

### Statistical analysis

Statistical analysis was carried out using PRISM 6 (GraphPad). Mann–Whitney test (comparing two groups), Kruskal–Wallis test (when compared to a single group), and Gehan–Breslow– Wilcoxon test (for survival curve comparison) were used in this study as indicated in the figure legends. Individual p values are indicated in the figure, with no data points excluded from statistical analysis. One representative experiment of at least three independent repeats is shown. For qPCR, each gene was normalized to a reference gene and compared with the Holm-Sidak test (individual comparison with paired value), where each p-value and DF were adjusted for respective multiple comparisons.

## Results

### miR-99 overexpression inhibits neutrophil motility and chemotaxis

We use the *lyzc* promoter to drive gene expression specifically in zebrafish neutrophils [26; 27]. The microRNA gene was inserted into the intron of a fluorescent reporter gene (fig.1A) to allow visualization of neutrophils expressing the transgene. To further characterize the zebrafish line *Tg(lyzC: miR-99-1/dendra2*)*^pu26^* with this referred to as miR-99, we first isolated neutrophils and performed miRNA RT-qPCR to confirm the miR-99 overexpression. Compared with the control line that only expresses the vector, the miR-99 level is significantly higher in the miR-99 line (fig. 1B). The two abundant neutrophil miRNAs, miR-223 and let-7e, were used as controls to demonstrate that this overexpression did not disrupt physiological microRNA levels. miR-99 overexpression did not affect the total neutrophil numbers in embryos (fig.1C, D) but reduced neutrophil recruitment to either the tail injury or ear infection sites (fig.1 E-H). In addition, the spontaneous neutrophil motility in the head mesenchyme was significantly slower upon miR-99 overexpression (fig 1I, J and Movie 1). We then sought to determine whether miR-99 inhibits human neutrophil migration. Here we used HL-60 cells, a myeloid leukemia cell line that can be differentiated into neutrophil-like cells as our model [25; 28]. We have developed a platform to express microRNAs in HL-60 cells after cell differentiation (fig. 2A) [25]. As expected, MIR-99 overexpression did not affect cell differentiation, cell death, or the expression of two other miRNAs, MIR-223 and LET-7 (fig. 2B-F). The differentiated (dHL-60) cells were then subjected to a transwell assay using N-Formylmethionyl-leucyl-phenylalanine (fMLP) as the chemoattractant. Overexpressing MIR-99, but not the vector control, significantly reduced chemotaxis (fig. 2G), indicating that overexpression of MIR-99 inhibits neutrophil migration in both zebrafish and human cell models. In addition, MIR-99 suppressed other functions in HL-60 cells, including phagocytosis, production of reactive oxygen species, and neutrophil extracellular traps (fig. 2H-J).

**Figure 1.**
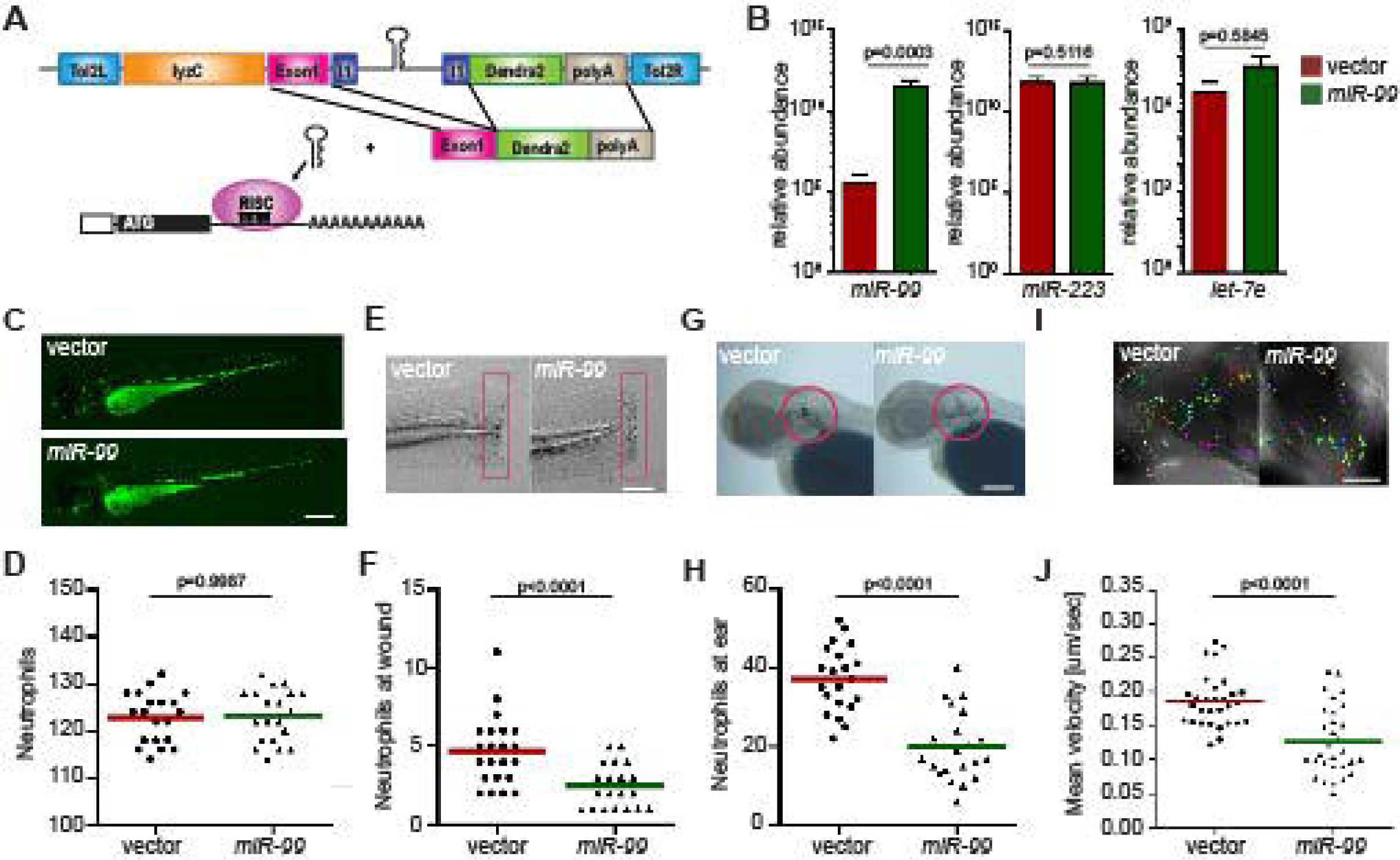
Neutrophil-specific *miR-99* overexpression hinders neutrophil recruitment and motility. (A) Schematic of the Tol2-lyzC:*miR-99*/Dendra2 plasmid, injected into wild-type AB zebrafish embryos to generate the transgenic line *Tg(lyzC:miR-99/Dendra2)* (*miR-99*) and *Tg(lyzC:Dendra2).* (B) The relative expression level of miR-99, miR-223, and let-7e (normalized to *U6* expression) in vector and miR-99 lines determined by RT-qPCR, mean ± s.d. (N = 3 biological replicates with ten larvae at each time point in each group). (C) and (D) Representative images and quantification of total neutrophils in vector and miR-99 lines. Scale bar: 500 μm. (E) and (F) Representative images and quantification of neutrophil recruitment to tail transection site in vector or miR-99 larvae at 1-hour post-injury. Neutrophils in the boxed region were quantified. Scale bar: 100 μm. (G) and (H) Representative images and quantification of neutrophil recruitment to localized ear infection in vector or miR-99 larvae at 1-hour post- infection. The infected ear is indicated with the circle. Scale bar: 100 μm (I) and (J) Tracks and quantification of neutrophil motility in vector or miR-99 larvae. Scale bar: 50 μm. All experiments were performed with F2 larvae at three dpf. One representative result of three independent experiments was shown (n=20). *P*-value was calculated with unpaired Student’s t- test.

**Figure 2.**
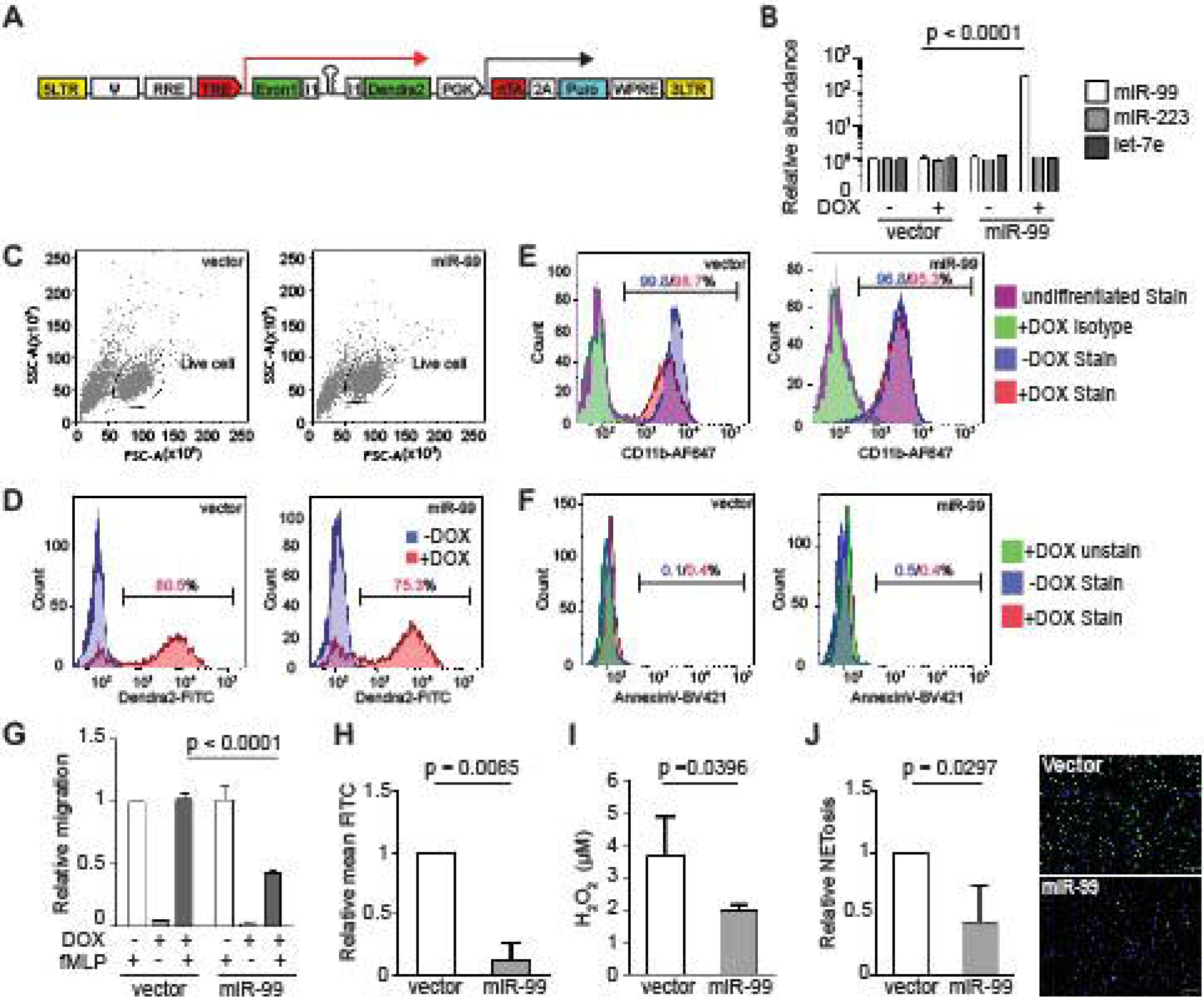
MIR-99 overexpression inhibits chemotaxis of dHL-60 cells. (A) Construct for inducible miRNA expression. The miRNA and a dendra2 reporter (green) are controlled by TRE, Tetracyclin response element (red). The PGK promoter drives constitutive expression of puromycin resistance gene (cyan) and the rtTA, reverse tetracycline-controlled transactivator (red). (B) Cell populations gated for downstream cell profile analysis and transwell quantification. (C) Doxycycline-mediated induction in the vector or miR-99 expressing dHL-60 lines. Cells without doxycycline-mediated induction were used as a baseline. Percentage of cells with dendra2 levels above the baseline are shown. (D) Surface CD11b levels in vector or miR-99 expressing dHL-60 cells. Undifferentiated cells or differentiated cells stained with an isotype control were used as a baseline. Percentage of cells with CD11b levels above the baseline with or without doxycycline-mediated induction are shown. (E) Annexin V staining in the total cell population (excluding debris) of vector or miR-99 expressing dHL-60 cells. An unstained live sample is used to determine the baseline. Percentage of cells with AnnexinV levels above the baseline with or without doxycycline-mediated induction are shown. (B-E) One representative experiment of three independent trials is shown. (F) Quantification of MIR-99, MIR-233, and LET-7E in dHL-60 cell lines with/without induced expression of the vector control or MIR-99. The result from three independent experiments is normalized to *U6* expression levels and presented as mean ± s.d., students’ t-test. (G - J) (G) Transwell migration of dHL-60 cells with/without induced expression of the vector control or MIR-99 toward fMLP, (H) phagocytosis of BioParticles by dHL60 cells represented as mean FITC, (I) H_2_O_2_ concentration in PMA-induced dHL60 cells, and (J) Quantification of dHL60 undergoing NETosis (left) and representative images (right). Scale bar, 100µm. Results are presented as mean ± s.d., from three independent experiments and normalized to vector, Kruskal–Wallis test or Mann-Whitney test.

### miR-99 suppresses the expression of RAR-related orphan receptor alpha RORα

To understand the underlying molecular mechanism of how miR-99 regulates neutrophil migration, we sequenced the transcriptome of neutrophils expressing miR-99. Since microRNAs generally suppress the expression of target transcripts [29], we focused on the down-regulated differentially expressed genes (DEG), which are associated with pathways in the cell cycle, microtubule-based movement, and DNA damage stimulus (fig. 3A, B and SI Appendix, Dataset S1, 2). Only two genes among the down-regulated DEGs, *roraa,* and *dec*1, contain predicted miR-99 binding sites in the 3’UTR (fig. 3C). Roraa is one of the retinoic acid receptor-related orphan receptor (ROR) gene family members in zebrafish. Identical to humans, there are three ROR genes in zebrafish: *rora*, *rorb,* and *rorc*. The protein product of *rora*, RORα, functions in metabolic regulation, energy homeostasis, thymopoiesis, and integration of the circadian clock[30; 31]. The functions of RORβ are mainly unknown, perhaps acts as a novel regulator in osteogenesis[32; 33]; RORγt, the protein product of *rorc*, guides Th17 cell differentiation and promoting IL-17 production[34]. Compared to *dec1*, the *roraa* gene orthologues are more conserved between zebrafish and humans (zebrafish *roraa* sequence is 89.96 % identical to its human ortholog *rora* while *dec1* in zebrafish is 59.55 % identical to its human orthologue). In addition, RORα polymorphisms correlate with different diseases, such as multiple sclerosis, breast cancer, asthma in human patients^56–58^. However, whether RORα plays a regulatory role in neutrophil migration is unknown. Therefore, we chose *roraa* to follow up. To validate the RNAseq results, we selected four other genes, *grn1, fabp7, tgfb1, il1b,* and confirmed their reduced transcript levels in the miR-99*-*overexpressing neutrophils (fig. 3D). To be noted, the expression of *roraa*, but not its homolog, *rorc*, was suppressed by miR-99. We then performed dual-luciferase reporter assays to determine whether miR-99 can directly bind to *roraa* 3’UTR and suppress reporter expression. miR-99 significantly reduced the relative luciferase activity, dependent on the *roraa* 3’UTRs (fig. 3E, F), indicating that miR-99 can directly target *roraa* in zebrafish.

**Figure 3.**
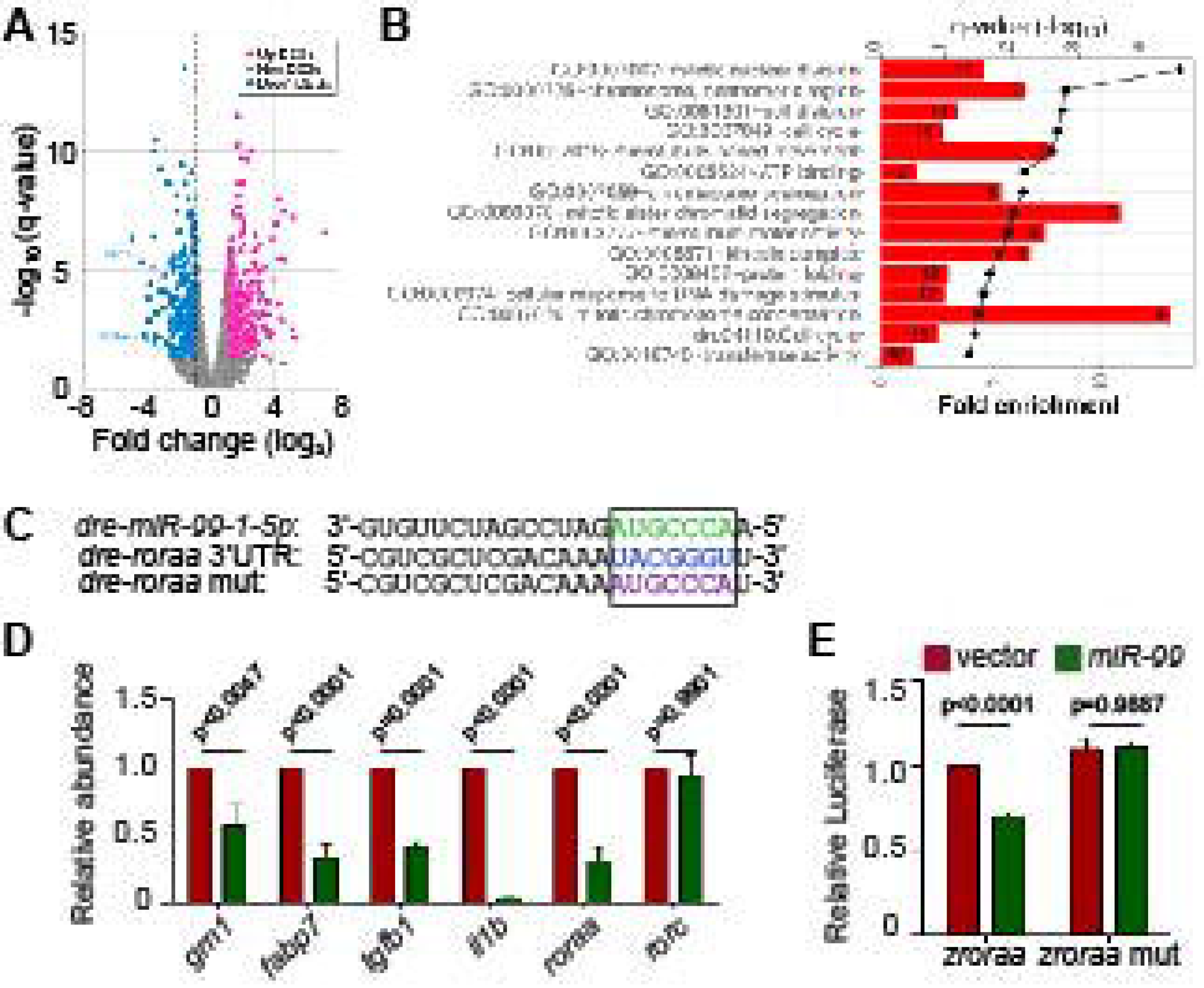
*miR-99* overexpression suppresses the expression of *roraa* in zebrafish neutrophils. (A) Volcano blot of DEGs with significant expression changes in miR-99 expressing neutrophils. Blue: down-regulated differentially expressed genes (DEGs); Magenta: up-regulated DEGs. (B) Significantly altered pathways in the down-regulated DEGs. (C) Alignment of miR-99-1-5p and predicted miR-99 binding sites in zebrafish *roraa* 3’UTR. (D) Transcript level quantification of predicted downregulated genes in neutrophils sorted from the vector or miR-99 zebrafish line. Results from three independent experiments are normalized to *rpl32* and presented as mean. Holm-Sidak test. (E) Suppression of Renilla luciferase expression via binding to zebrafish *roraa* 3’UTRs by miR-99. Results from three independent experiments are normalized to firefly luciferase and presented as mean ± s.d., Kruskal–Wallis test.

### Pharmacological inhibition of rora decreases neutrophil chemotaxis in both zebrafish and human

RORα binds DNA as a monomer and constitutively activates transcription [35; 36]. Although the endogenous ligands of this receptor are not entirely clear, synthetic ligands can be used to suppress or enhance RORα activities [37]. To test whether Rorα regulates neutrophil migration in homeostasis and inflammation conditions, we used SR3335, an inverse agonist of RORα, SR2211, a RORγt inverse agonist, and VPR66, an inverse agonist of both RORα and RORγt.

None of the inhibitors affected zebrafish survival or neutrophil numbers (fig. S1A). Neutrophil motility was significantly explicitly reduced by the RORα, but not the RORγ inhibition (fig. 4A, B and Movie 2). Consistently, neutrophil recruitment to the infected ear (fig. 4C, D) or tailfin amputation sites (fig. 4E, F) was significantly and specifically inhibited by the RORα inhibitor. Human neutrophils possess limited transcriptional activity and do not require transcription for chemotaxis [38; 39]. To ensure that our findings are not restricted to zebrafish, we isolated primary human neutrophils and treated them with the ROR inverse agonists. Similar to our observation made in zebrafish, the inverse agonist of RORα, but not RORγ, reduced chemotaxis of primary human neutrophils 1 hr after treatment, without causing apoptosis (fig. 4G, H, fig.

**Figure 4.**
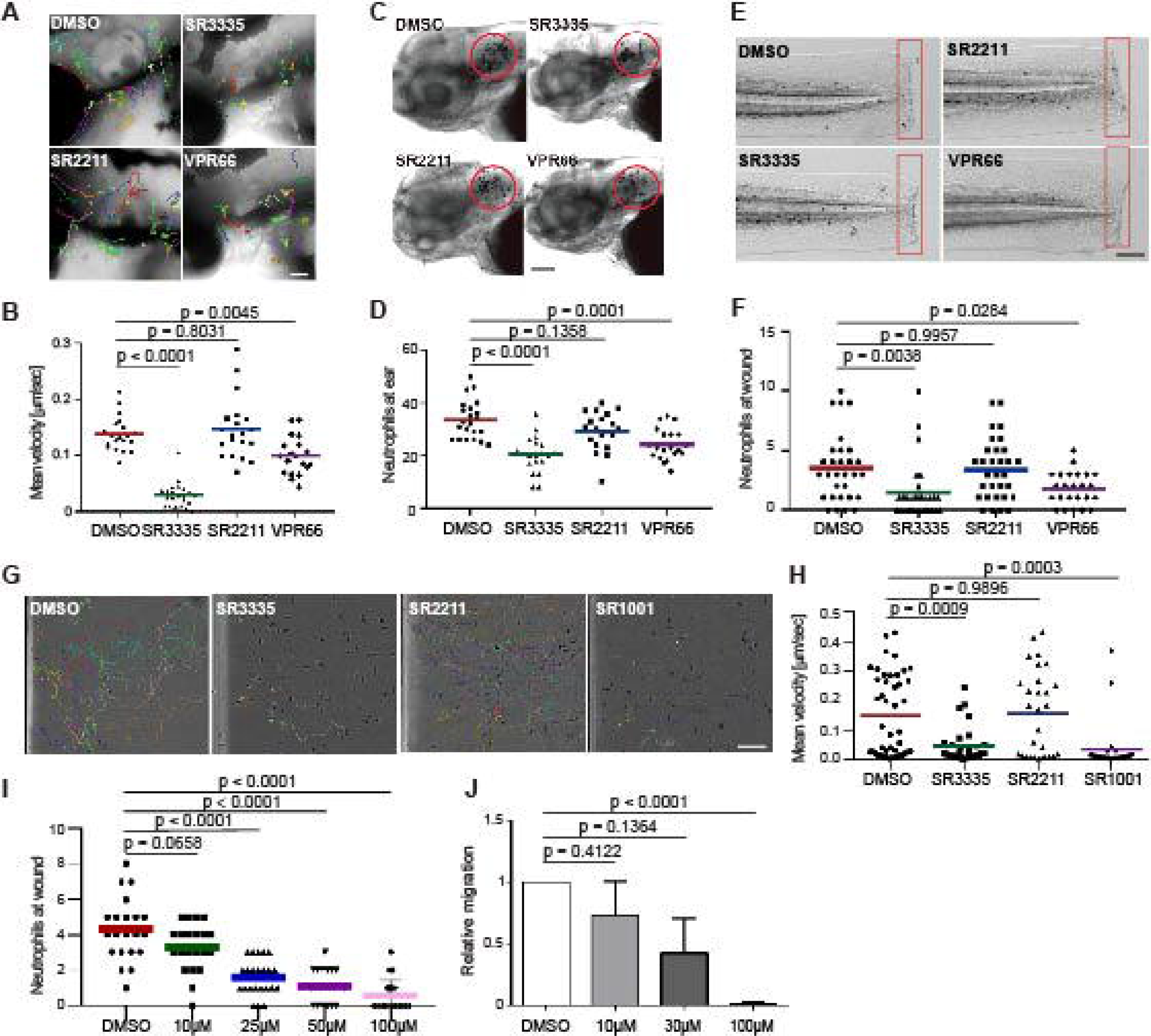
Pharmacological inhibition of Rora reduces neutrophil motility and chemotaxis in zebrafish and humans. (A, B) Tracks and quantification of neutrophil motility in zebrafish larva treated with SR3335 (100μM), SR2211 (100μM), or VPR66 (25μM). Scale bar, 200 µm. Three embryos, each from three different founders, were imaged. Quantification of neutrophils in one representative video is shown, Kruskal–Wallis test. (C, D) Representative images and quantification of neutrophils recruited to the infected ear in zebrafish larva treated with RORα specific inhibitor (SR3335, 100μM), RORγ specific inhibitor (SR2211, 100μM) or pan-ROR family inhibitor (VPR66, 25μM). Scale bar, 100 µm. (E, F) Representative images and quantification of neutrophils recruited to tail fin transection sites in zebrafish larva treated with SR3335 (100μM), SR2211 (100μM), or VPR66 (25μM). Scale bar: 500 µm. (C-F) The result from one representative experiment is shown as mean, Mann–Whitney test. (G, H) Representative tracks and mean velocity of primary human neutrophils treated with SR3335 (50μM), SR2211 (50μM), or SR1001 (pan-ROR family inhibitor) (50 µM) migrating towards fMLP in 3D matrigel. Scale bar, 100 µm. Representative results for three individual trials are shown. The result is presented as mean, Mann–Whitney test. (I) Neutrophil recruitments after zebrafish tail wounding at different dosages of SR3335 treatment compared to 1% DMSO treatment, Kruskal–Wallis test. (J) Transwell migration of primary human neutrophils treated with DMSO (0.1%) or SR3335 at 10, 30, or 100 μM toward 100 nM fMLP. Results are presented as mean ± s.d., from three independent experiments and normalized to DMSO (0.1%), Kruskal–Wallis test.

S1B-G and Movie 3). In addition, the ROR inverse agonists SR3335 suppressed neutrophil chemotaxis in fish and humans in a dose-dependent manner (fig4 I, J). In summary, we report a previously unrecognized role for RORα in regulating neutrophil chemotaxis.

### DNA binding activity of Rora is required for neutrophil motility

Due to the potential off-target effects of the inhibitors, we next sought genetic evidence for RORα function in neutrophil migration. RORα protein consists of one DNA-binding domain and one ligand-binding domain, working as a monomer and activates transcription when binding to the DNA response element. A truncated version of RORα lacking the ligand-binding domain acts as a dominant-negative by occupying the DNA response elements without activating transcription. We constructed zebrafish lines overexpressing Rorα (1-180aa) in neutrophils, *Tg(lyzC:mcherry-2a-rora-dn)^pu29^*, together with a control line *Tg(lyzC:mcherry-2a)^pu30^*(fig.5A). Despite multiple attempts, we cannot generate the line overexpressing the full-length *rora* as a control. Expressing the *rora* DN did not affect total neutrophil numbers but reduced neutrophil motility in the head mesenchyme (fig.5 B-E and Movie 4). Consistently, Rorα DN significantly reduced neutrophil recruitment to infection or tail amputation sites (fig. 5F-I). The control or the Rorα DN line was crossed with *Tg(mpx: GFP-UtrCH*) to label stable actin in neutrophils [19]. In the control neutrophils, stable actin was enriched in the retracing tail. In comparison, in Rorα DN expressing neutrophils, stable actin mislocalized to cell front or the middle of the cell body, indicating disrupted cell polarity (fig.5J and Movie 5). In addition, we knocked down *rora* specifically in neutrophils using the CRISPR-Cas9 system recently optimized in our lab where Cas9 expression is restricted in neutrophils and two sgRNAs targeting *rora* are expressed ubiquitously [40]. Consistently, neutrophil migration is significantly inhibited upon *rora* deletion (fig.5K-N and Movie 6). Taken together, our results indicate that the RORα mediated transcription is required for neutrophil polarization and efficient chemotaxis.

**Figure 5.**
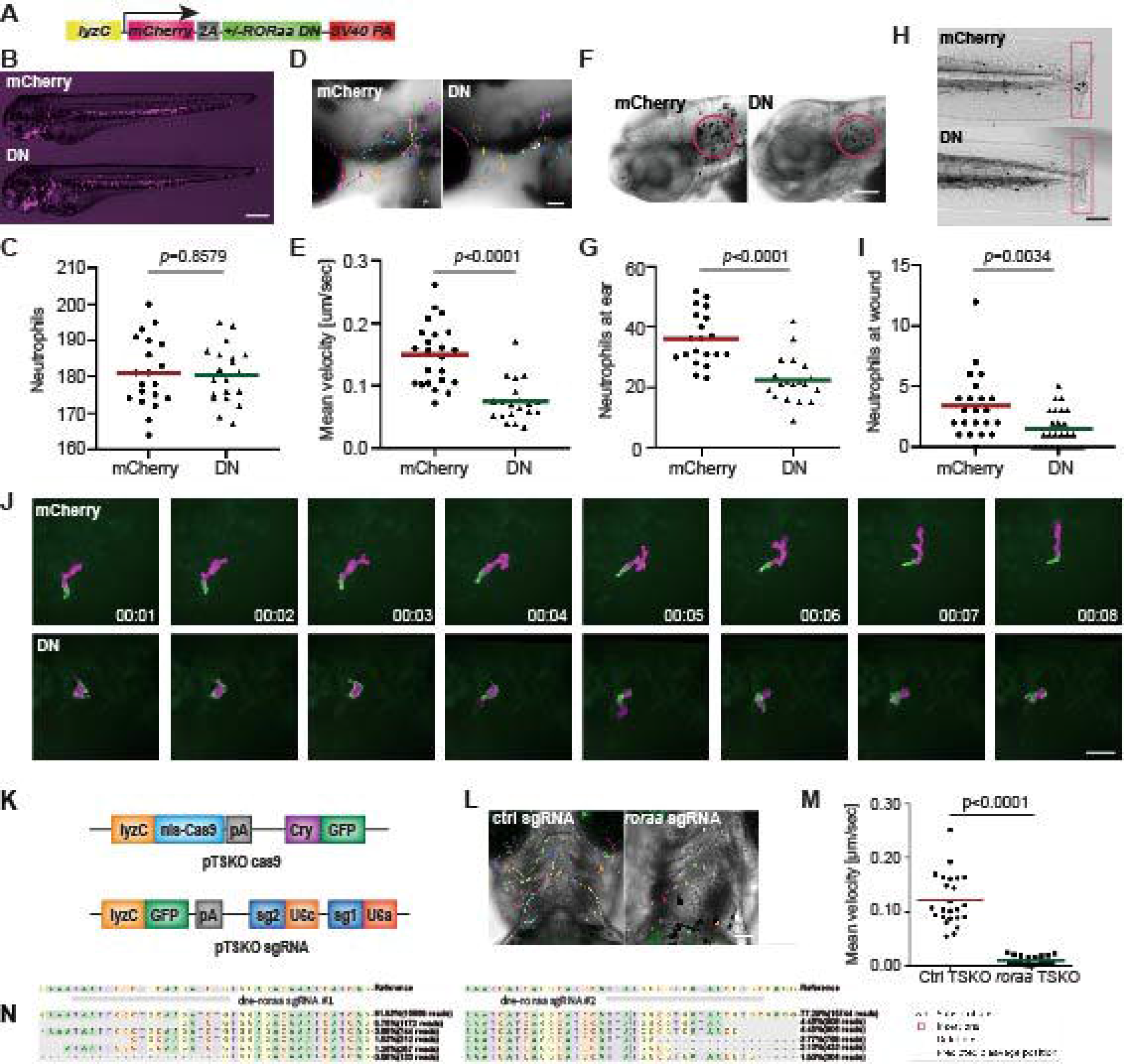
Dominant-negative Rorα suppresses neutrophil motility and chemotaxis. (A) Schematic of the construct for neutrophil-specific expression of vector control or *ror*α dominant-negative (DN). (B, C) Representative images and quantification of total neutrophil numbers in vector or *ror*α DN zebrafish line. Scale bar: 500 μm. (D, E) Representative images and velocity of random neutrophil migration in vector or *ror*α DN zebrafish line. Scale bar, 100 µm. Three embryos each from three different founders were imaged, and quantification of neutrophils in one representative video is shown, Kruskal–Wallis test. (F, G) Representative images and quantification of neutrophils recruited to the infected ear in vector or *ror*α DN zebrafish line. Scale bar, 100 µm. (H, I) Representative images and quantification of neutrophil recruitment to the tailfin transection sites in vector or *ror*α DN zebrafish line. Scale bar, 200 µm. The result is presented as mean, Mann–Whitney test. (J) Simultaneous imaging of utrophin-GFP distribution in neutrophils expressing either mCherry alone or with Rora DN. Data are representative of more than three separate time-lapse videos. Scale Bar, 50 µm. (B, C, F-I) Representative results for three individual trials are shown. (K) Schematics of the plasmid constructs for neutrophil-specific Cas9 expression (upper construct) and for ubiquitous expression of sgRNAs with a neutrophil-specific expression of GFP (lower construct). (L, M) Representative images (L) and quantification (M) of neutrophil motility at head region of 3dpf larvae from *Tg(LyzC: Cas9, Cry: GFP, U6a/c: rora guides, LyzC: GFP)* fish. Student’s t-test. Scale bar, 100 µm. (N) Deep sequencing of the rora loci targeted by guide RNAs in (K). The sequences on the top are wild-type sequences, and the five most frequent types of mutations are shown. Point mutations, deletions, and insertions are all observed.

### RORα is required for defense against bacterial infection

Our current finding suggests that miR-99 overexpression or RORα inhibition suppresses neutrophil chemotaxis. We then investigated whether RORα is required for defending against bacterial infection using *P. aeruginosa* [41]. Survival of the zebrafish embryos was significantly reduced upon expression of miR-99 or Rorα DN in neutrophils (fig. 6A, B). In line with this observation, inhibiting RORα, but not RORγ, using small molecules also reduced survival (fig. 6C, D). Taken together, RORα plays an essential role in regulating neutrophil activation and the defense against bacterial infection.

**Figure 6:**
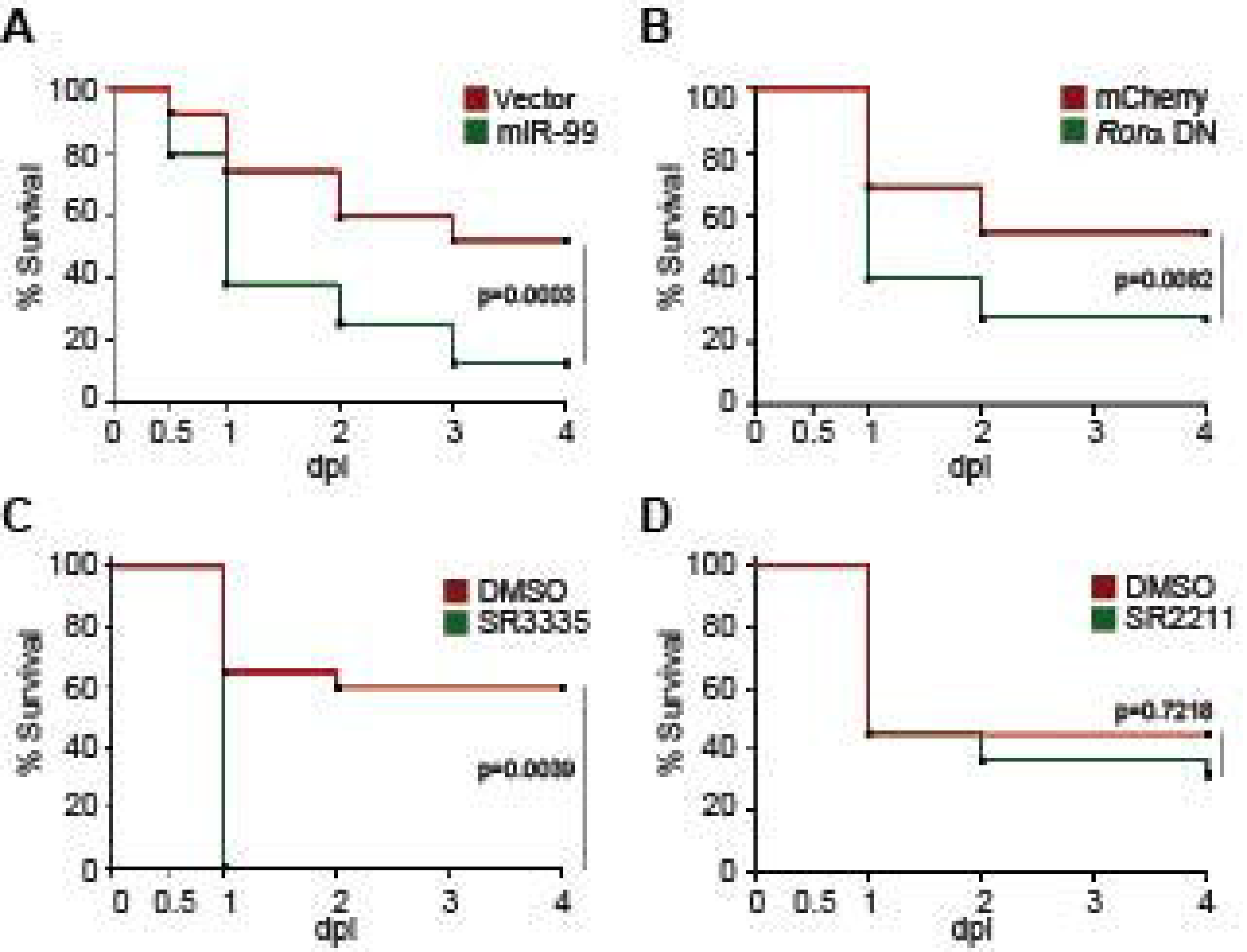
Rorα in neutrophils protects zebrafish against *Pseudomonas aeruginosa* infection. (A) Survival curve of 3 dpf larvae from the vector or *miR-99* line after i.v. infection with *Pseudomonas aeruginosa* (PAK). (B) Survival curve of 3 dpf larvae from the mcherry or *roraa* DN line larvae after i.v. infection with PAK. (C) Survival curve of 3 dpf larvae treated with DMSO or SR3335 (100µM) injected with PAK. (D) Survival curve of 3 dpf larvae treated with DMSO or SR2211 (100µM) injected with PAK. One representative experiment of three independent biological repeats (n = 20 each group) is shown. The result is analyzed with Gehan– Breslow–Wilcoxon test.

### Rorα regulates receptor signaling pathways in zebrafish neutrophils

RORα is expressed in primary human and zebrafish neutrophils, but not in the human neutrophil model cell line, HL-60, after different differentiation methods [42]. Due to the possible off-target effects of inhibitors and the difficulty of genetically altering primary human neutrophils, we used zebrafish neutrophils overexpressing the Rorα DN to understand the RORα regulated gene network., 216 DEGs are down-regulated in zebrafish neutrophils overexpressing Rorα DN compared with the control with a cut off of false discovery rate of 0.05. However, the consensus ROR binding motif (AxxTxGGTCA) is not enriched in the downregulated DEGs, suggesting direct and indirect transcriptional regulation by RORα. The downregulated DEGs are enriched in the cell death, transmembrane signaling receptor activity, and protein phosphorylation pathways. Receptor signaling and protein phosphorylation pathways are involved in cell migration and activation, supporting the biological outcome of RORα inhibition in neutrophils. We further compared our gene list to a previously published list of RORα target genes in monocytic and endothelial cells identified using chromatin immunoprecipitation (ChIP) [43]. We converted downregulated DEGs to their human homologs, among which, ten are putative RORα target genes in both THP-1 and HUVEC cells, and 13 and 23 are putative RORα target genes in either one of the cells (Table 1). Finally, we selected five transcripts that are putative RORα target genes to validate this RNA seq result (fig. 7D). As expected, the level of *roraa* is significantly higher due to overexpression of the DN construct. Rho GTPase Activating Protein 17 (Arhgap17b) is a signaling molecule regulating Cdc42 and Rac1 [44]. Protein Tyrosine Kinase 2 Beta (Ptk2ba) regulates protein tyrosine kinase activity and is implicated in human neutrophil migration [45] and host defense response to bacterial infection [46]. MHC Class I Polypeptide- Related Sequence B (Mhc1zba) is essential in self-antigen presentation under stressed conditions recognized by innate-like lymphocytes [47]. Calcium Voltage-Gated Channel Auxiliary Subunit Gamma 3 (Cacng3b) is among the most significantly altered DEGs. It is a voltage-dependent calcium channel subunit on the plasma membrane, regulating the receptor activity [48]. Another most significantly changed DEG we validated is the transcription elongation regulator one like (Tcerg1l). Together, the validated downregulation of these molecules provides a molecular understanding of how RORα-dependent gene expression is required for proper neutrophil migration and activation.

**Figure 7:**
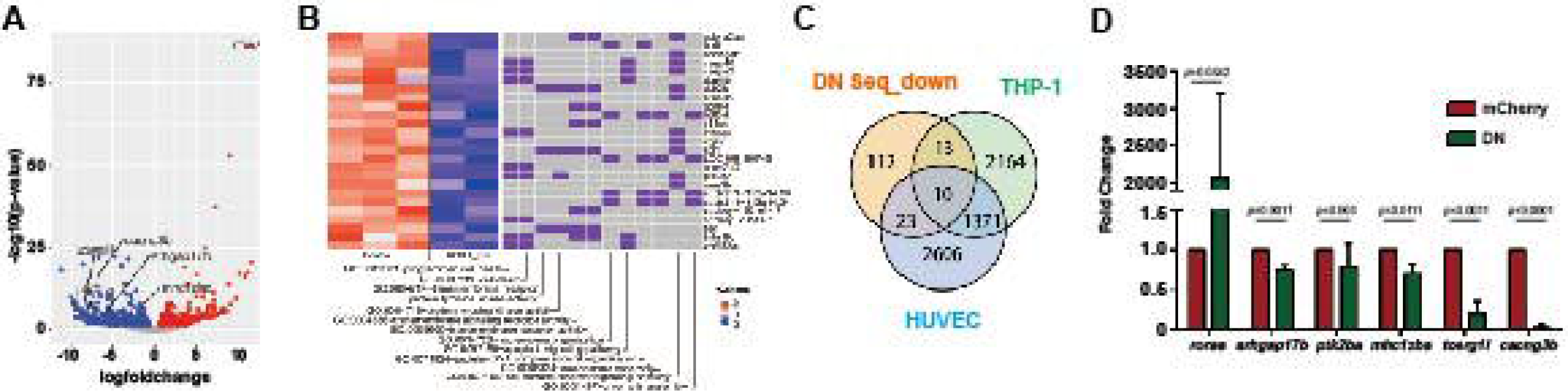
RNA-Seq reveals the direction of *ror*α possible downstream genes in neutrophil migration. (A) Volcano blot of DEGs with significant expression changes in *ror*α DN neutrophils. Cyan: down-regulated differentially expressed genes (DEGs); Magenta: up-regulated DEGs. (B) Significant gene function enrichment of human homologs of genes in the down-regulated DEGs. The heat map indicates the log2 fold change of the selected DEGs. The relevance of individual DEGs in each pathway is indicated with purple. (C) Venn diagram showing human homologs of genes in the down-regulated DEGs overlapping with putative RORα target genes in THP-1 or HUVEC cells. (D) Transcript level validation of downregulated genes in neutrophils sorted from *ror*α DN line zebrafish. Results from three independent experiments are normalized to *rpl32* and presented as mean ± s.d., students’ t-test.

## Discussion

In the present study, we characterized miR-99 as a critical regulator of neutrophil migration and chemotaxis. miR-99 hinders neutrophil motility by downregulating genes enriched in zebrafish receptor activity pathways and directly targeting *roraa*. We further demonstrate that the transcriptional regulation activity of RORα is critical for neutrophil migration and defense against a bacterial pathogen.

MIR-99 has multiple regulatory roles in various biological processes, especially hematopoietic and leukemic stem cell differentiation and function [49]. The miR-99/100∼125 microRNA cluster block of the transforming growth factor β (TGFβ) pathway enhances Wnt signaling to promote the survival and expansion of hemotopoeitc stem cells megakaryocytic differentiation [50]. miR-99 is highly expressed in hematopoietic stem cells (HSCs) and acute myeloid leukemia stem cells (LSCs) and promotes their self-renewal and leukemia-initiating activity through targeting Hoxa1 [51]. Outside the blood system, miR-99 is a tumor suppressor. miR-99 suppresses prostate cancer cell proliferation by suppressing the expression of prostate-specific antigen [52]. It also stops expanding the human cervical cancer cell line, HeLa, by targeting Tribbles 2 [53]. MiR-99 suppresses FGFR3 expression in lung cancer and Erk1/2 and Akt, reducing cell growth and metastasis [54]. In addition, miR-99 is transiently induced after radiation, reducing the efficiency of DNA repair by suppressing SNF2H expression [55]. miR-99 family is highly downregulated in physiological but up-regulated during pathological hypertrophy [56]. No prior studies have investigated the role of miR-99 in neutrophils.

Therefore, our discovery that miR-99 is a suppressor of cell migration in leukocytes expands the biological functions of miR-99 outside cancer biology.

We identified that retinoid acid receptor-related orphan receptor α (RORα) is a direct target of miR-99. RORα regulates embryonic development, metabolism, circadian rhythm, and inflammation [57; 58; 59]. Dominant-acting RORα mutations are associated with autism and cerebellar ataxia in humans [60]. RORα polymorphisms correlate with human diseases such as breast cancer [61] and asthma [62]. RORα gene expression is downregulated in the blood of multiple sclerosis patients [63]. Recent studies suggested prominent role of RORα in regulating T cell and ILC2 development [64; 65] and immune cell functions, including T regulatory cells in allergic skin inflammation [66], Th17 driven inflammatory disorders [34; 67], survival of CD4+ T cells in colitis [68] and lung infection [69], ant-tumor immunity of liver-resident natural killer cells/ILC1s [70], macrophage activation in LPS-induced septic shock [71] and diet-induced obesity [72], and ILC3 driven fibrosis [73]. Both pro-inflammatory [74; 75] and anti- inflammatory [76; 77] roles of RORα are observed in different cell types. RORα deficient mast cells and macrophages produce increased amounts of TNF-alpha and IL-6 upon activation [76]. Ectopic expression of ROR alpha1 inhibits TNF alpha-induced IL-6, IL-8, and COX-2 expression in human primary smooth-muscle cells, through the induction of a significant inhibitory protein I kappa B alpha [77]. In contrast, RORα suppressed the expression of suppressors of cytokine signaling 3 (SOCS3), a negative regulator of inflammation, and deletion of RORα protects against oxygen-induced proliferative retinopathy in a murine model [74].

RORα stimulates inflammation in adipose tissue by enhancing the expression of ER stress response genes and inflammatory cytokines [75]. In the intestinal epithelial cell, RORα attenuates NF-kB activity and inhibits excessive inflammation in a mouse colitis model [78]. In human neutrophils, endotoxin administration downregulates the expression of RORα [79].

However, it is not previously known if RORα plays a role in regulating neutrophil function.

Although we characterized the gene expression network regulated by RORα in zebrafish neutrophils and provided the molecular understanding that RORα regulates receptor activation and downstream signaling pathways, the limitation of the current study is that the results are yet to be confirmed in human neutrophils. The most surprising result is that chemotaxis of primary human neutrophils is reduced after only 1 hr of RORα inhibitor treatment, given that human neutrophils possess limited transcriptional activity and do not require transcription for chemotaxis [38; 39]. One possibility is that SR3335 has off-target effects, and the phenotype is not related to RORα. Another scenario is that RORα activity needs to be maintained to support neutrophil chemotaxis. The previous work identified limited transcriptional activity [39] used differentiated HL-60 cells, which does not express RORα. Therefore, the conclusion should be revisited in primary human neutrophils. However, due to the short half-life of primary human neutrophils, metabolic labeling to identify nascent mRNA in this cell type is challenging. A list of active transcripts in zebrafish neutrophils is also obtained by isolating associated ribosomal mRNAs [80]. Although this method can identify genes responsive to tissue damage, due to the high background in the IP (false positives), this result cannot be used to infer a reliable, complete list of active transcripts. Future studies will be needed to carefully set up the assays to get to the functional transcripts in human neutrophils and identify those regulated by RORα. In addition, neutrophils display rhythmical behavior mediated by core clock genes in mice [81] and zebrafish [82]. In the future, it is also interesting to determine how RORα regulates the rhythmical behavior of neutrophils.

In summary, our data indicated that the miR-99/RORα axis functions as a novel modulator in the acute state of neutrophil recruitment and is potentially a novel target in preventing and treating acute inflammatory diseases and infections.

## Authorship Contributions

AH, SM, CS, JW, and DQ designed research and wrote the manuscript. AH, TW, RS, WZ, JW, ZC, ST, ZC performed the experiments. AH, TW, RS, KL, and SL analyzed data. All authors read and approved the manuscript.

## Data Availability Statement

The RNA-seq raw and processed data are submitted to the Gene Expression Omnibus (GSE144873, GSE171501). Plasmids are available on Addgene.

