## Supplemental Figures for "RORA regulates neutrophil migration and activation in zebrafish"

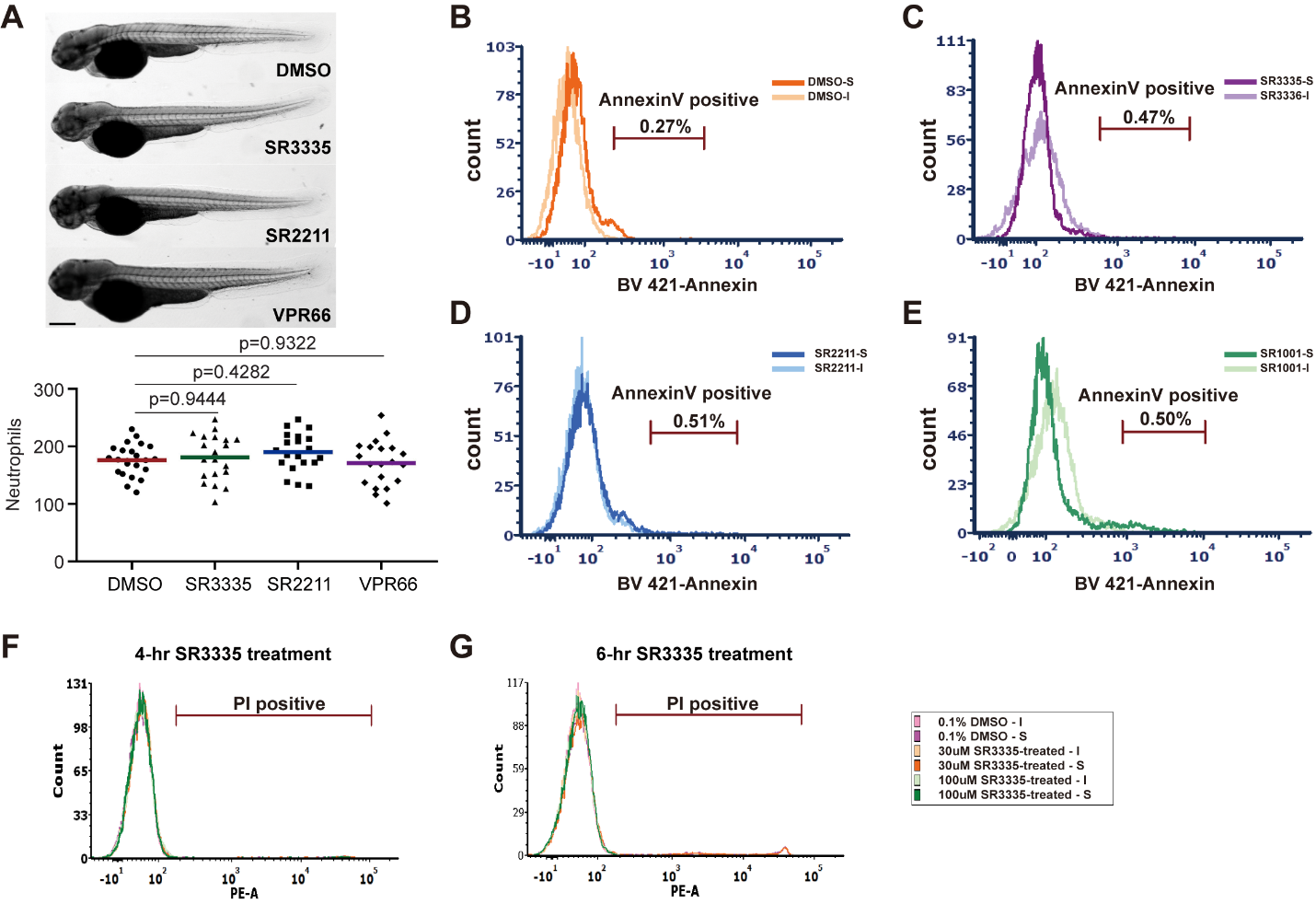

**Figure S1. RORα inhibitor treatment does not cause neutrophil death.** (A) Representative images and quantification of total neutrophils in zebrafish larva treated with RORα specific inhibitor (SR3335, 100μM), RORγ specific inhibitor (SR2211, 100μM) or pan-ROR family inhibitor (VPR66, 25μM). *P*-value was calculated with unpaired Student's t-test. Scale bar: 100 µm. (B-E) Annexin V staining in the total cell population (excluding debris) of primary human neutrophils treated with SR3335 (50μM), SR2211 (50μM), or SR1001 (pan-ROR family inhibitor) (50 µM). Percentage of cells with Annexin V levels above the baseline are shown. (F-G) PI staining for neutrophils treated with SR3335 for 4 or 6 hours. S: signal; I: unstained cells.

**Movie S1: Tracked movies of neutrophil motility lines in the head mesenchyme of the vector and the *miR-99*.**

The video shows the motility of neutrophils in 3 dpf vector and miR-99 zebrafish lines. Videos were recorded for 30 min with a 1 min interval. Representative videos from n = 3 independent experiments with three fish in each group are shown. Scale bar: 100 µm.

**Movie S2: Tracked movies of neutrophil motility in the head mesenchyme treated with vehicle, SR3335, SR2211, or VPR66.**

The video shows the motility of neutrophils in 3 dpf zebrafish larvae treated with DMSO (vehicle control), SR3335, SR2211, or VPR66. Videos were recorded for 30 min with a 1 min interval. Representative videos from n = 3 independent experiments with three fish in each group are shown. Scale bar: 100 µm.

**Movie S3: SR3335 inhibits chemotaxis of primary human neutrophils.**

Tracks of primary human neutrophils chemotaxis treated with DMSO, SR3335, SR2211, or SR1001 towards fMLP. Videos were recorded for 50 min with 30 s intervals. Representative videos from n = 4 separate trials are shown. Scale bar: 50 µm.

**Movie S4: Tracked movies of neutrophil motility in the head mesenchyme of the control and Rorα DN lines.**

The video shows the motility of neutrophils in 3 dpf control or Rorα DN zebrafish lines. Videos were recorded for 30 min with 1 min interval. Representative videos from n = 3 independent experiments with three fish in each group are shown. Scale bar: 100 µm.

**Movie S5: Morphology of neutrophils from the control and Rorα DN lines**

The video shows the morphology and stable actin distribution of neutrophils in 3 dpf control or Rorα DN zebrafish lines crossed with a *Tg(mpx:gfp-utrCH)* line. Videos were recorded for 10min with a 1min interval. Representative videos from n = 3 independent experiments with three fish in each group are shown. Scale bar: 20 µm.

**Movie S6: Tracked movies of neutrophil motility in the TSKO lines.**

The video shows the motility of neutrophils in 3 dpf larvae from the control and *roraa* neutrophil-specific knock-out fish lines. Videos were recorded for 30 min with a 1 min interval. Representative videos from n = 3 independent experiments with three fish in each group are shown. Scale bar: 100 µm.

**Dataset S1. List of genes that are downregulated in the miR-99 overexpressing neutrophils.**

**Dataset S2. Pathway enrichment analysis of downregulated genes in *miR-99* overexpressing neutrophils.**

**Dataset S3. List of genes that are downregulated in the Roraa-DN overexpressing neutrophils.**

**Dataset S4. Pathway enrichment analysis of downregulated genes in Roraa-DN overexpressing neutrophils.**

**Table 1. Overlap human homologs of genes in the down-regulated DEGs with putative RORα target genes in THP-1 or HUVEC cells.** Genes selected for validation in Figure 7d are highlighted.
